# Constructing a full-length single-nucleus transcriptomic atlas of the obese mouse by UURNA-seq

**DOI:** 10.64898/2026.07.14.738381

**Authors:** Xiaoping Han, Xueyi Wang, Lei Yang, Lifeng Ma, Hanyu Wu, Renying Wang, Guoji Guo

**Author notes:** These authors contributed equally (Xueyi Wang, Lei Yang, Lifeng Ma).

## Abstract

Single-nucleus RNA-seq (snRNA-seq) offers advantages in sample preparation for atlas construction and data mining of gene regulation. However, current snRNA-seq protocols struggle to balance sensitivity and throughput. Here we present ultra-throughput and ultra-sensitivity single-nucleus total RNA sequencing (UURNA-seq), a platform that enables large-scale atlas construction in a single day. UURNA-seq outperforms existing single-cell and single-nucleus protocols, with reduced coverage bias, higher gene detection efficiency, and greater throughput at lower per-cell cost. This performance extends to primary tissues and is consistent in fresh and frozen samples, supporting flexible workflows for clinical and archival specimens. Notably, UURNA-seq also detects substantially more non-coding RNAs, including lncRNAs and sncRNAs, alongside mRNAs at single-nucleus resolution, uncovering previously inaccessible layers of cellular regulation in complex tissues. Leveraging these capabilities, we constructed the first comprehensive multi-tissue atlas of obesity and characterized RNA dynamics at atlas scale, revealing both conserved and tissue-specific transcriptional alterations and identifying critical dysregulation of hormone signaling and metabolic pathways. Together, these results establish UURNA-seq as a sensitive, scalable, and cost-effective platform that provides a route to dissect the molecular basis of complex diseases such as obesity.

## Introduction

In the past decade, single-cell RNA sequencing (scRNA-seq) has revolutionized our understanding of cellular heterogeneity in both healthy and diseased states^1^. However, the isolation of high-quality single-cell suspensions remains challenging, particularly for tissues that are difficult to dissociate, and harsh enzymatic dissociation may introduce cell type bias. To address these limitations, single-nucleus RNA sequencing (snRNA-seq) has emerged as a viable alternative that is compatible with frozen or fixed tissues^2,3^. Current methods for single-cell resolution include droplet-based microfluidic systems^4^, microwell-based platforms^4,5^, and split-pool strategies^6–8^. While these approaches have substantially enhanced the throughput of sc/snRNA-seq, advancing beyond earlier low-throughput technologies^9–11^, library construction cost and limits on per-experiment throughput still restrict their broader use. Furthermore, comprehensive transcriptome coverage also remains challenging for current high-throughput methods.

Most sc/snRNA sequencing technologies rely on barcoded oligo-dT primers hybridizing to the poly(A) tail of transcripts for RNA capture and subsequent cDNA synthesis. However, this approach results in significant 3’ end bias, compromising full-length transcript information. Moreover, intracellular non-polyadenylated transcripts, including certain non-coding RNAs, are not detected. Recent advances in full-length transcriptome sequencing have enabled the detection of non-coding and non-polyadenylated RNAs, with sequencing reads spanning the entire gene body^12–16^. However, these methods, whether plate-based^12,13,15,16^ or droplet-based^13,14^, are constrained by their dependence on sorting instruments or microfluidic systems.

Obesity is a major risk factor for many metabolic disorders, including type 2 diabetes, cardiovascular diseases, and certain cancers^17^. Obesity also strongly affects endocrine function, disrupting hormonal balance and inter-tissue communication^18^. This endocrine disruption shows clear sexual dimorphism, particularly in women, who are more susceptible to obesity-related hormonal perturbations^19^. Notably, estrogen signaling plays a central role in energy homeostasis and adipose tissue distribution, as shown by the increased visceral adiposity in postmenopausal women^20^. The impact extends to reproductive function, where obesity disrupts the hypothalamic-pituitary-ovarian axis, leading to complications including anovulation, menstrual irregularities, and infertility^21^. Despite these systemic effects, our understanding of obesity’s impact on inter-tissue hormonal networks at the single-cell level remains limited.

To address these challenges, we developed UURNA-seq, a full-length single-nucleus RNA-seq platform that combines *in situ* reverse transcription with random primers and four rounds of split-pool barcoding, enabling ultra-high-throughput profiling without specialized instrumentation. Benchmarking UURNA-seq against established platforms showed markedly reduced coverage bias, higher gene detection efficiency, and lower per-cell cost. We further demonstrated its robustness in primary tissues and in both fresh and frozen samples, supporting flexible workflows for clinical and archival specimens. Applying UURNA-seq to the adult mouse brain, we generated a comprehensive single-nucleus atlas encompassing 42 cell types, and found that UURNA-seq captured neuronal transcriptomes with substantially reduced dissociation-induced stress signatures relative to 10× Chromium. Notably, UURNA-seq detected more non-coding RNAs, particularly lncRNAs and sncRNAs, with higher sensitivity than the 10× snRNA-seq platform. Using its ultra-high throughput, we applied UURNA-seq to build a batch-effect-free multi-tissue obesity atlas comparing wild-type and ob/ob mice, encompassing 749,804 single nuclei from 14 tissues enriched for neuronal and endocrine systems. This atlas enabled systematic profiling of cell-type composition, differential gene expression, metabolic and hormonal pathway remodeling, and transcription factor regulatory networks associated with obesity. Together, these results establish UURNA-seq as a sensitive, cost-effective, and scalable tool for full-length single-nucleus transcriptomics and discovery of non-coding RNA.

## Results

### Design and workflow of UURNA-seq

The UURNA-seq protocol began with the isolation of nuclei followed by fixation using 4% paraformaldehyde (PFA) to facilitate in situ reverse transcription (RT). Fixed nuclei were distributed into 96-well plates and reverse-transcribed with barcoded random primers using multiple annealing and looping-based amplification cycles (MALBAC)^22^, introducing the first barcode (Fig. 1a). These random primers hybridized efficiently to internal transcript regions rather than only to the poly(A) tail, thereby capturing non-polyadenylated RNAs in addition to mRNAs. After RT, nuclei were pooled and residual primers were removed by Exonuclease I digestion, and *in situ* poly(A) tailing was then added to the 3’ end of the newly synthesized cDNA. A second round of split-pool barcoding was performed using oligo-dT primers carrying the next barcode, and excess primers were blocked by a 3’ dideoxy-C modification to prevent barcode contamination. After pooling and a third splitting, the resulting RNA–DNA hybrids were treated with RNase and subjected to bidirectional strand extension by DNA polymerase. Nuclei were then lysed with low-concentration SDS, and after SDS neutralization with Tween-20 the released transcripts underwent PCR pre-amplification to incorporate the third barcode. Finally, the sequencing library was constructed directly with primers bearing the P5 and i7 adapters, with the i7 primer serves as the fourth barcode (Fig. 1b). The complete library structure is shown in Fig. 1c.

**Fig. 1.**
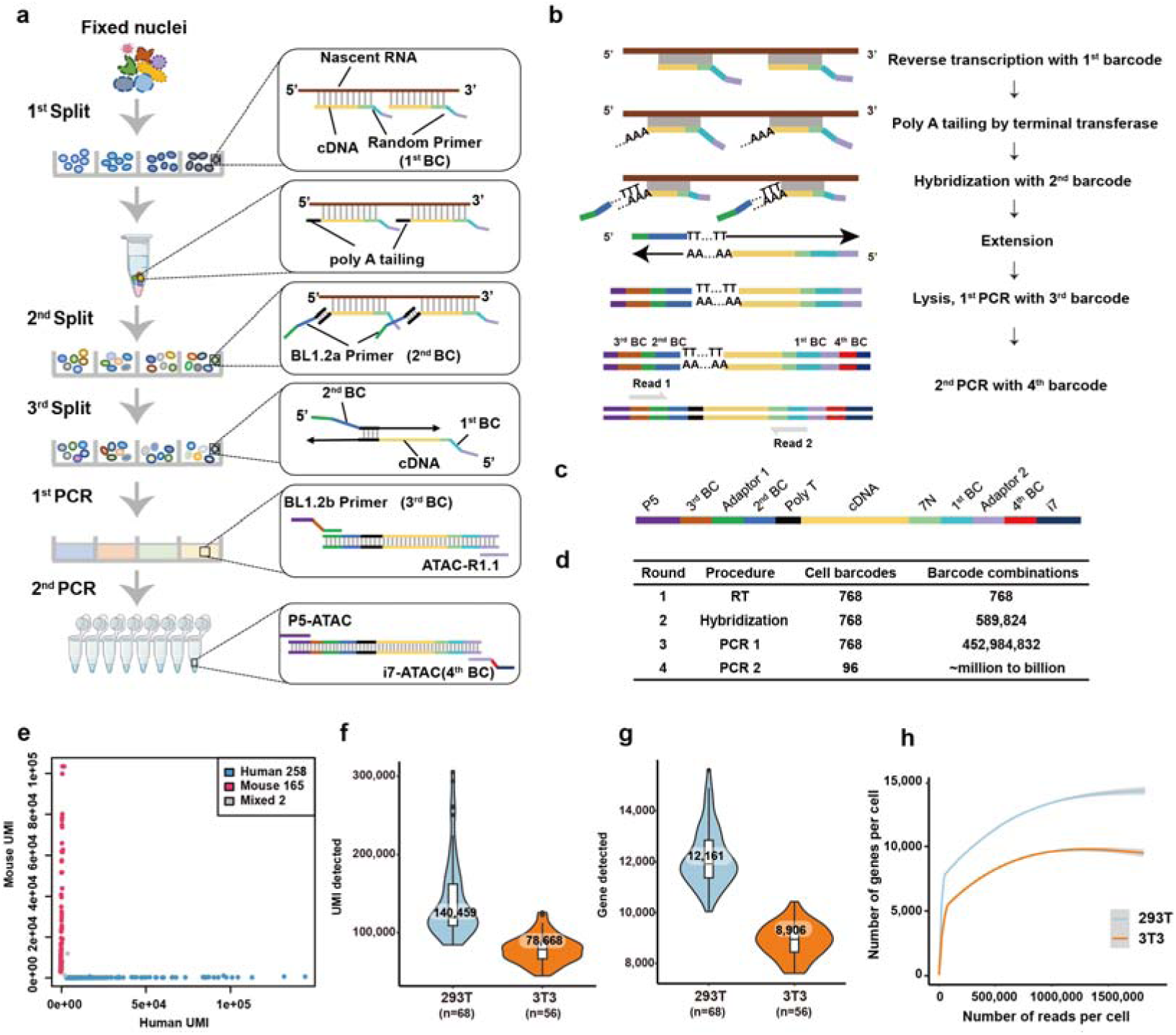
UURNA-seq workflow and performance. **a**, Overview of the UURNA-seq workflow. Single nuclei are extracted and fixed, then subjected to reverse transcription (RT) with barcoded random primers. Nuclei are pooled, the cDNA 3’ ends are polyadenylated, and the nuclei are split again and hybridized with barcoded primers. Nuclei are then pooled and split again for second-strand synthesis, followed by in-well lysis. The cDNA is amplified by PCR with barcoded primers, purified, and amplified in a second PCR to generate the library. **b,** Structure of the cDNA intermediate at each step of UURNA-seq library construction. **c,** Schematic representation of the library structure for UURNA-seq. **d,** Number of barcodes used in each round of split-pool and the resulting number of barcode combinations. **e,** Collision rate in a human-mouse species-mixing experiment, using HEK 293T (blue) and NIH/3T3 (red) cells. Barcodes with more than 20% of detected UMIs from the other species were classified as doublets/mixed (grey). **f, g,** Detection sensitivity of UURNA-seq, shown as UMI counts (f) and gene counts (g). Only cells with a sequencing depth of at least 50,000 reads were included (HEK 293T, n = 68; NIH/3T3, n = 56). **h,** Number of genes detected per cell as a function of reads per cell, for HEK 293T and NIH/3T3 cells. Gene detection approaches saturation well below the maximum depth sequenced, so high sensitivity is achieved at relatively shallow depth.

These four rounds of split-pool barcoding provide two key advantages. First, successive rounds of combinatorial labeling expand the barcode space multiplicatively (Fig. 1d, Supplementary Table 1), allowing multiple samples to be labeled and processed in a single run, reducing batch effects. UURNA-seq therefore achieves ultra-high throughput, reaching millions of cells per experiment. Second, because barcoding relies solely on split-pool operations, the method requires no microfluidic or sorting instrumentation and is therefore simpler and more accessible.

We next validated the data quality of UURNA-seq. A species-mixing experiment with mouse NIH/3T3 and human HEK 293T cells confirmed its single-cell resolution, yielding a low doublet rate of 0.47% (Fig. 1e). We next assessed its sensitivity by deep sequencing, detecting a median of 140,459 UMIs and 12,161 genes per 293T nucleus and 78,668 UMIs and 8,906 genes per 3T3 nucleus (Fig. 1f, g). Gene detection rose steeply at low depth and reached a clear inflection point at approximately 100,000 reads per cell (Fig. 1h), indicating high sensitivity.

### Performance and robustness of UURNA-seq

UURNA-seq employs random primers, which hybridize efficiently to any region of a transcript and thereby recover full-length transcriptome data. To test this capability, we benchmarked UURNA-seq against established single-cell and single-nucleus platforms in 293T cells. Unlike platforms exhibiting pronounced 5’ or 3’ end bias (10× Chromium^23,24^ and sci-RNA-seq^6^), UURNA-seq produced a full-length coverage profile that closely resembled that of established full-length sequencing platforms (Smart-seq3^12^, FLASH-seq^24^) (Fig. 2a). Building on this coverage advantage, we next assessed gene detection efficiency across sequencing depths. At 5,000–30,000 reads per cell, UURNA-seq detected more genes per cell on average than any other platforms evaluated (Fig. 2b), and this advantage persisted at saturation (approximately 200,000 raw reads per cell), where it detected significantly more genes than sci-RNA-seq^6^ and SPLiT-seq^8^ (Fig. 2c). Together, these results demonstrate that UURNA-seq achieves greater detection sensitivity and makes more efficient use of sequencing reads. Beyond its high sensitivity and independence from specialized instrumentation, UURNA-seq offers a markedly lower per-cell cost than comparable platforms while still enabling ultra-high throughput (Fig. 2d).

We next evaluated whether these advantages extend to primary tissue, in which cellular heterogeneity and dissociation pose greater challenges. In the adult mouse pituitary gland, UURNA-seq resolved seven distinct endocrine cell clusters from 273 nuclei, with median counts of 6,000–8,000 genes per nucleus (Fig. 2e–f), confirming that the sensitivity is preserved in complex primary tissue.

**Fig. 2.**
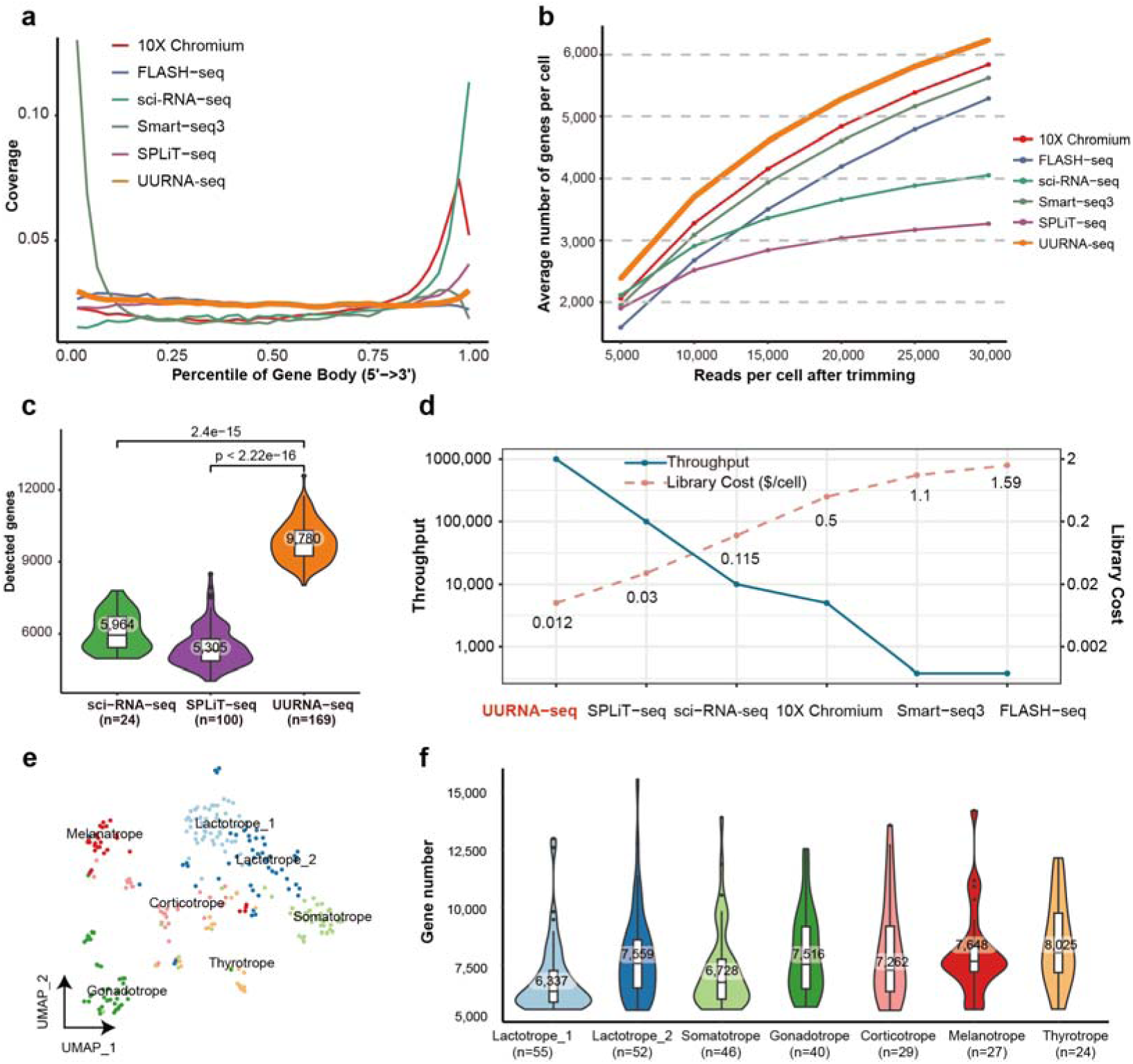
Benchmarking UURNA-seq against existing single-nucleus RNA-seq methods. **a**, Comparison of gene body coverage along the gene across different platforms. **b,** Gene detection sensitivity versus sequencing depth in HEK 293T cells, showing the number of detected genes plotted against quality-filtered reads per cell at varying downsampling thresholds. Only cells with a sequencing depth of at least 30,000 reads were included (UURNA-seq: n = 664, FLASH-seq: n = 100, Smart-seq3: n = 100, SPLiT-seq = 277, 10× Chromium: n = 3,512, sci-RNA-seq: n = 43). **c,** Gene detection sensitivity of UURNA-seq compared with major snRNA-seq methods in single HEK293T nuclei. Only cells with a sequencing depth of at least 200,000 reads were included (UURNA-seq, n = 169; SPLiT-seq, n = 100; sci-RNA-seq, n = 24). **d,** Library cost and throughput of UURNA-seq compared with other methods. Source data for the other RNA-seq methods is available in a systematic comparison study. UURNA-seq exhibited the highest throughput per experiment and the lowest cost per cell. **e,** UMAP visualization of the adult mouse pituitary. **f,** Number of detected genes across annotated cell types corresponding to (e). UURNA-seq also showed high sensitivity in adult tissues.

However, samples are often more complex and are frequently frozen rather than fresh, particularly in clinical settings. We therefore evaluated the compatibility of UURNA-seq with cryopreservation, and its performance with different surfactants, to determine whether it maintains cellular integrity and representative sampling under these conditions. To assess cryopreservation, we stored post-poly(A)-tailed nuclei at –80°C for one or two days and processed them alongside fresh nuclei (Fig. 3a–b). Cryopreserved and fresh samples showed minimal batch effects by UMAP (Fig. 3c) and high concordance in gene expression (R = 0.99) (Fig. 3d), with only marginally higher nuclei recovery from fresh samples (Fig. 3e), indicating compatibility with frozen tissue and potential for clinical application.

**Fig. 3.**
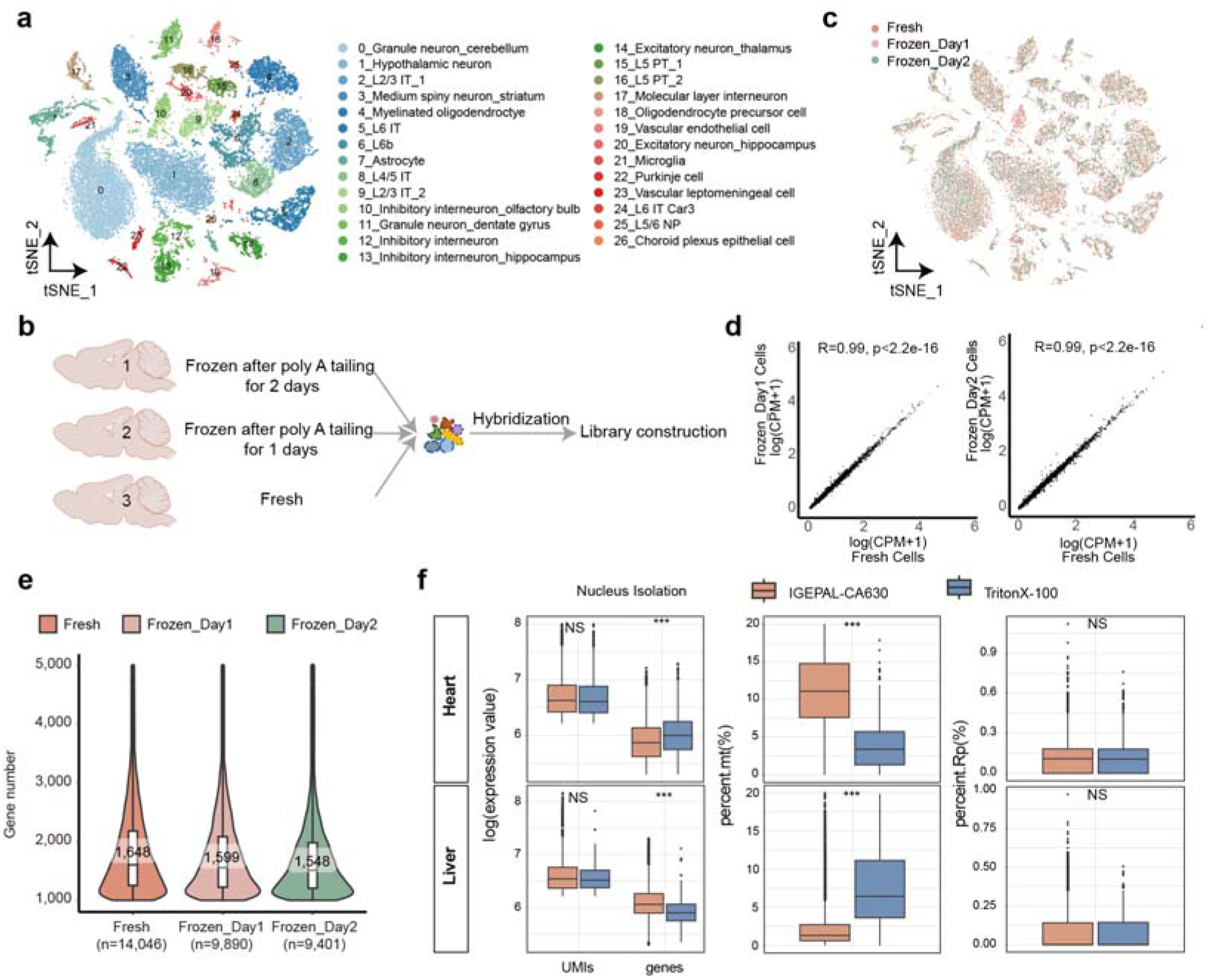
Benchmarking UURNA-seq across fresh and frozen tissues. **a**, t-SNE visualization of quality-controlled cells from fresh and frozen tissues, colored by annotated cell types. **b,** Flowchart of the experimental design. After poly(A) tailing, hybridization was performed alongside fresh brain, followed by library construction. **c,** t-SNE visualization of quality-controlled cells from fresh and frozen tissues, colored by samples. **d,** Correlation between gene expression measurements in UURNA-seq profiles of fresh nuclei vs. frozen_day1 nuclei (left) and fresh nuclei vs. frozen_day2 nuclei (right). Fresh and cryopreserved tissues correlated closely (R = 0.99, p < 2.2 × 10^-16^). **e,** Violin plots of detected genes per cell across fresh and cryopreserved samples, with medians indicated. Sensitivity remained stable in both fresh and frozen tissues. **f,** Box plots comparing log-transformed expression values (left), the percentage of mitochondrial (mt) genes (middle), and the percentage of reads mapped to rRNA (rp) (right) for heart and liver using two nucleus isolation buffers: IGEPAL-CA630 (orange) and TritonX-100 (blue). Box plots: centre line, median; boxes, first and third quartiles of the distribution; whiskers, highest and lowest data points within 1.5 × IQR.

**Fig. 4.**
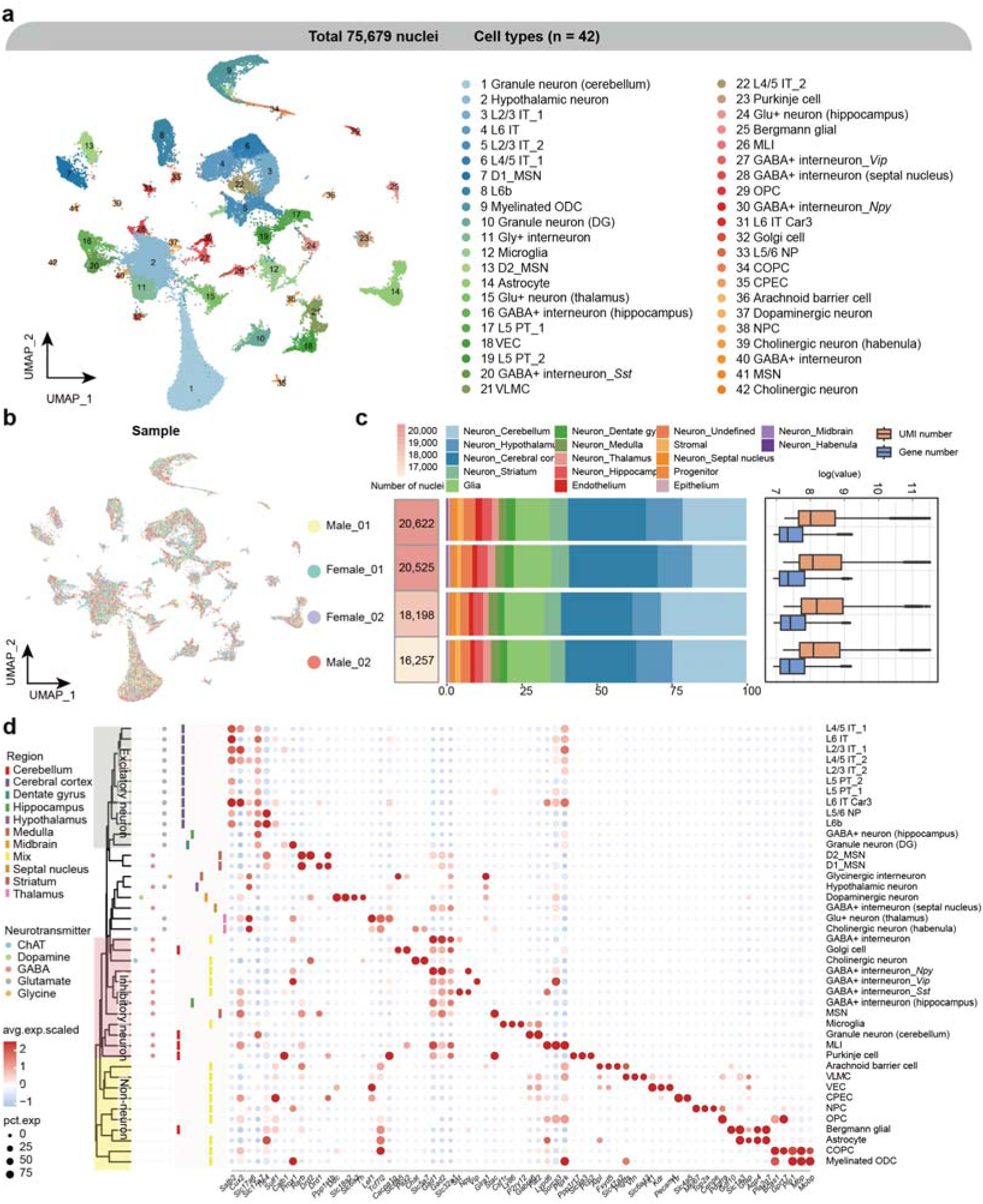
UURNA-seq atlas of the adult mouse brain. **a**, UMAP visualization of 75,679 nuclei after quality control and annotation, colored by cell types. D1-MSN, D1-type medium spiny neuron; D2-MSN, D2-type medium spiny neuron; ODC, oligodendrocyte; OPC, oligodendrocyte precursor cell; COPC, committed oligodendrocyte precursor cell; NPC, neural progenitor cell; CEPC, choroid plexus epithelial cell; DG, dentate gyrus; MLI, molecular layer interneuron; VEC, vascular endothelial cell; VLMC, vascular leptomeningeal cell. **b,** UMAP visualization of single-nucleus transcriptomes from four mouse brain samples (two males, two females). **c,** Stacked bar chart of the proportional distribution of lineages and neuronal regions, with total nuclei counts indicated. Box plots show the distribution of UMI and gene counts detected per nucleus across samples. **d,** Dot plot of cell-type-specific gene expression across neuronal and non-neuronal populations. Dot size represents the percentage of cells expressing each gene (pct.exp) and color intensity indicates the scaled average expression (avg.exp.scaled). Left annotations indicate brain regions and neurotransmitter types. Cell types are hierarchically clustered and include major neuronal subtypes (excitatory and inhibitory neurons), glial cells (astrocytes and oligodendrocytes) and vascular cells.

Nuclear isolation is another critical determinant of snRNA-seq data quality^6–8,25^, especially for tissues poorly suited to direct grinding under liquid nitrogen. We therefore compared commonly used detergents across tissues and found that IGEPAL-CA630 yielded superior performance in liver, whereas Triton X-100 was more effective in heart, as assessed by gene detection rates and mitochondrial read fractions (Fig. 3f). Together, these results show that UURNA-seq is robust across diverse tissue types and preservation conditions, and that its workflow accommodates direct tissue freezing, nuclei storage or storage after poly(A) tailing prior to sequencing. Collectively, these findings position UURNA-seq as a high-sensitivity, low-cost, and scalable platform for full-length single-nucleus transcriptomics, with robustness that supports its broad application to primary and clinically relevant samples.

### Benchmarking with 10× Chromium using whole-brain scRNA-seq data

To generate a broadly applicable reference, we built a single-nucleus atlas of the adult mouse brain using UURNA-seq, profiling nuclei from two male and two female mice to capture biological variation across both sexes. UMAP analysis of 75,679 high-quality nuclei resolved 42 distinct neuronal and glial populations, all consistently represented across individual samples, indicating reproducible recovery of cell types rather than sample– or sex-specific artifacts (Fig. 3a–c). Region-specific markers enabled confident identification of individual populations, including striatal medium spiny neurons (*Ppp1r1b*, *Drd1*, *Drd2*^26^) and cerebellar granule neurons (*Gabra6*, *Fat2*^27^) (Fig. 3d; Supplementary Table 2).

We next compared UURNA-seq with 10× Chromium, one of the most widely used scRNA-seq platforms. To enable a fair comparison, we first matched analogous cell types between the two datasets using MetaNeighbor (correlation > 0.95), ensuring that any differences would reflect the profiling method rather than mismatched populations (Fig. 5a). Across matched populations, UURNA-seq exhibited significantly weaker dissociation-induced stress signatures (Fig. 5b, c). Consistent with this, differential expression analysis showed that genes enriched in UURNA-seq were primarily associated with core neuronal functions, including synapse regulation and axonogenesis, whereas downregulated genes were enriched for mitochondrial metabolism (Fig. 5d, e; Supplementary Table 3). The pattern extended across neuronal populations, which showed predominant upregulation in UURNA-seq (Fig. 5f). UURNA-seq detected higher expression of neuronal regulators (*Kcnq5, Kcnip4)* and glial development factors (*Npas3*^28^*, Mir99ahg*^29^), whereas 10× Chromium captured the elevated mitochondrial and ribosomal gene expression (*Cox8a*, *mt-Cytb*).

**Fig. 5.**
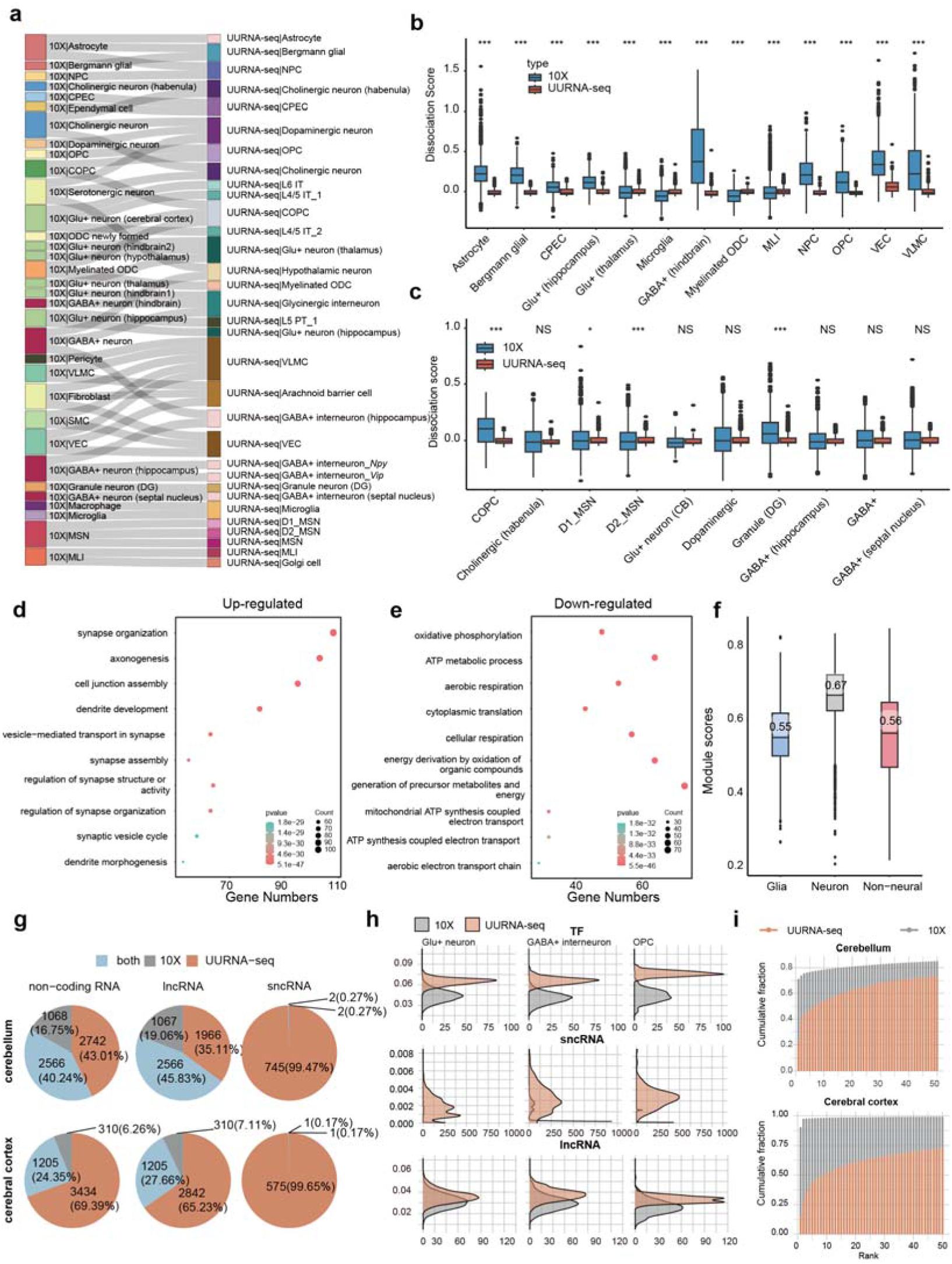
Differentially expressed genes in UURNA-seq mouse brain dataset compared with 10× Chromium single-cell dataset. **a**, Sankey diagram of the correspondence between cell types identified by 10× Chromium (left) and UURNA-seq (right), demonstrating the hierarchical relationships and concordance between cell type annotations from two platforms (UURNA-seq, 75,679 nuclei; 10× Chromium, 66,939 nuclei). **b, c,** Box plots of dissociation-related stress signature scores across brain cell types for UURNA-seq (brown) and 10× Chromium (blue). The y axis represents normalized expression scores calculated as background-corrected log (TP10K+1) values. Cell types are indicated on the x axis. Statistical significance was assessed using the Wilcoxon rank-sum test and corrected by the Benjamini-Hochberg method (***, FDR < 0.001; **, FDR < 0.01; *, FDR < 0.05; NS, not significant). Boxes indicate the interquartile range (IQR) with the median line; whiskers extend to 1.5 × IQR; points represent outliers. **d, e,** Gene Ontology (GO) pathways enriched among up-regulated DEGs (d) and down-regulated DEGs (e) in UURNA-seq relative to 10× Chromium (|logFC| > 1 and *P* < 0.01). Dot size represents the number of genes associated with each term and the color gradient indicates the statistical significance (*P* value) of the enrichment. **f,** Box plots of module scores for glial, neuronal, and non-neural modules of up-regulated genes. **g,** Pie charts comparing the detection of three categories of non-coding RNAs (total non-coding RNA, lncRNA and sncRNA). **h,** Density plots of the relative abundance of transcript biotypes in three typical neural cell types for 10× Chromium (gray) and UURNA-seq (brown): TF (top), sncRNAs (middle) and lncRNA (bottom). **i,** Cumulative UMI distribution of ranked lncRNA genes for 10× Chromium (grey) and UURNA-seq (brown) single-nucleus data from cerebellum (top) and cerebral cortex (bottom).

Since these differences largely stem from the distinct RNA compartments sampled, we next compared UURNA-seq with 10× Chromium 3’ v3 single-nucleus data, which profile the same nuclear compartment. In this matched comparison, UURNA-seq detected non-coding RNAs more effectively, particularly sncRNAs, which 10× largely missed (Fig. 5g). Biotype analysis across neuronal and glial populations showed that UURNA-seq detected a higher proportion of transcription factors (> 2-fold) and uniquely identified sncRNAs (Fig. 5h). This advantage was especially pronounced for lncRNAs. In the 10× data, *Malat1* alone accounted for over 80% of lncRNA counts owing to AT-content bias, whereas UURNA-seq detected lncRNAs far more broadly (Fig. 5i).

### Construction multi-organ transcriptomic atlas of obese mice by UURNA-seq

UURNA-seq’s ultra-high throughput enabled construction of a batch-effect-free multi-tissue atlas comparing wild-type (WT) and *ob/ob* mice, comprising 14 tissues, with an emphasis on neuronal and endocrine systems (Fig. 6a, Supplementary Table 4). After quality control, we derived expression profiles from 749,804 single nuclei (346,233 WT, 408,720 *ob/ob*). Clustering, visualized by UMAP (Fig. 6b), resolved 55 major clusters organized by lineage (Supplementary Fig. 1a). Gene expression profile was highly specific across different cell types (Supplementary Fig. 1b). Notably, the atlas included previously underreported endocrine tissues^5,25,30,31^, such as pituitary, adrenal and thyroid glands, which contributed to the secretory lineage characterization (Supplementary Fig. 1c). The stromal lineage (C22, C31 and C34) was derived predominantly from adipose tissues, aorta, and stromal cells. Co-expression analysis of the stromal lineage revealed enhanced stromal functionality in obese mice, evidenced by upregulation of fibroblast, smooth muscle cell-specific, and pan-stromal modules (Supplementary Fig. 2a, b). These modules were enriched for extracellular matrix organization and vascular-associated smooth muscle cell development, implicating stromal remodeling as a critical response to obesity^32,33^ (Supplementary Fig. 2c).

**Fig. 6.**
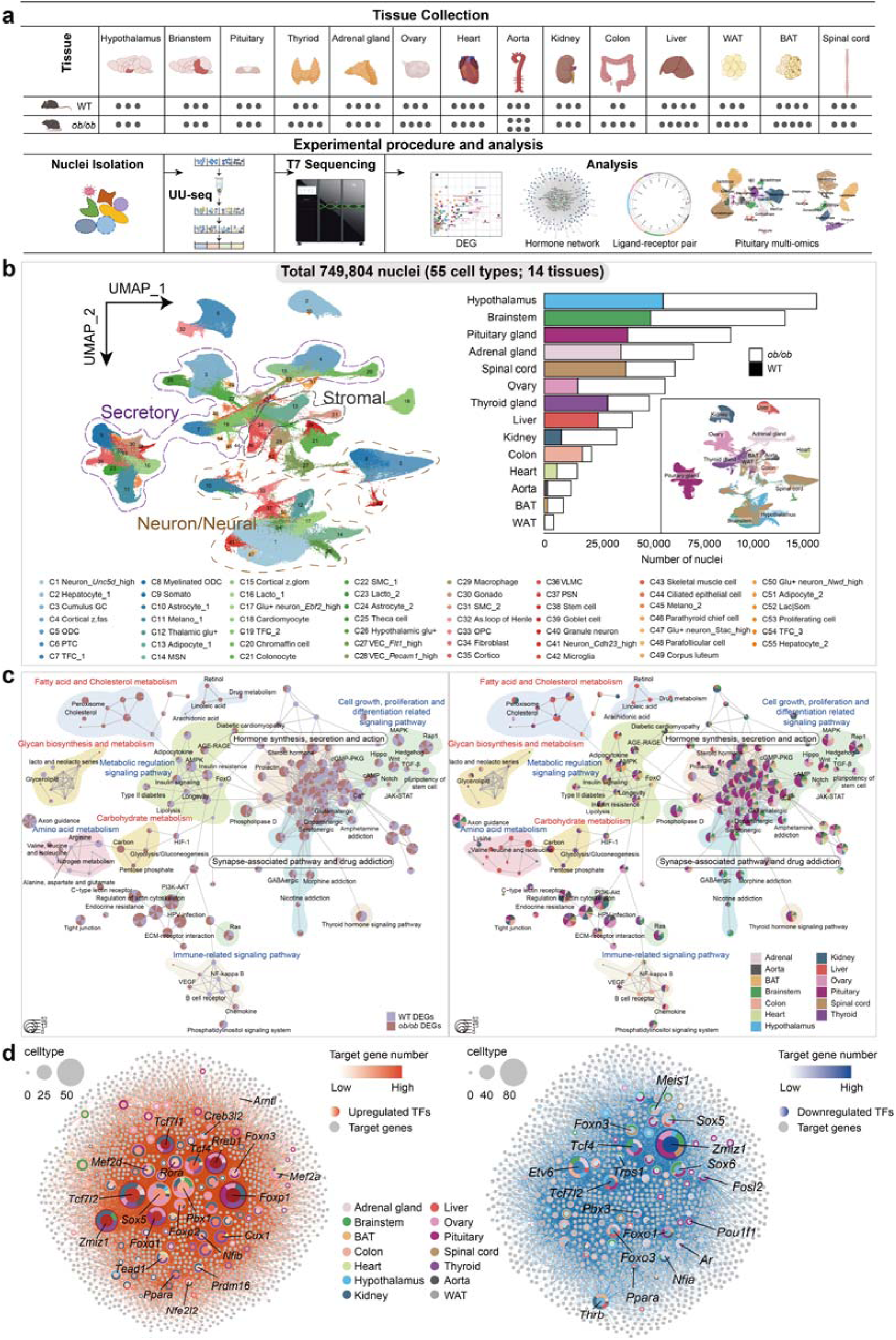
Cross-tissue snRNA-seq atlas of normal and obesity adult mice. **a**, Experimental design of data collection and analysis flow. A total of 14 tissues were collected, with at least two samples per tissue for each mouse type. **b,** Cross-tissue single-nucleus atlas. UMAP visualization of single-nucleus profiles (dots) colored by broad cell types (left). Major lineages are highlighted and circled with different colors. Bar chart of nuclei counts per tissue (right). UMAP visualization of the atlas colored by tissue was shown at the lower right. Cumulus GC, Cumulus granulosa cell; Cortical z.fas, Cortical zona fasciculata cell; ODC, Oligodendrocyte; PTC, Proximal tubule cell; TFC, Thyroid follicular cell; Somato, Somatotrope; Melano, Melanotrope; Thalamic glu+, Thalamic glutamatergic neuron; MSN, Medium spiny neuron; Cortical z.glom, Cortical zona glomerulosa cell; Lacto, Lactotrope; Glu+ neuron, Glutamatergic neuron; Smooth MC, Smooth muscle cell; Hypothalamic glu+, Hypothalamic glutamatergic neuron; VEC, Vascular endothelial cell; Gonado, Gonadotrope; As.loop of Henle, Ascending loop of Henle; OPC, oligodendrocyte precursor cell; Cortico, Corticotrope; VLMC, Vascular leptomeningeal cell; PSN, Peripheral sensory neuron; Skeletal MC, Skeletal muscle cell; Lac|Som, somalactotrope. **c,** Network plots of KEGG pathways enriched among the DEGs between lean and obese mice. The left panel represents the analysis categorized by mouse type, while the right panel presents the data stratified by tissue. **d,** Network plots of TFs up-regulated (left) and down-regulated (right) in obesity. Node size indicates the number of cell types. Node color, from light to dark, represents the number of target genes, from low to high.

Clustering of all tissues other than the pituitary gland (analyzed in detail below) is presented in Supplementary Figs.3–5, with tissue-specific markers listed in Supplementary Table 5. These figures show marker genes for each cell type and differential abundance between lean and obese mice (Supplementary Table 6). Differentially expressed genes (DEGs) revealed the most pronounced obesity-related differences in endocrine tissues and metabolic organs (Supplementary Fig. 6a). Augur analysis identified lactotropes, somatotropes, theca cells and granulosa cells as the cell types most responsive to obesity (Supplementary Fig. 6b). Additionally, CELLECT analysis showed strong associations between BMI and hormone-secreting cells in the pituitary and specific neuronal subtypes, underscoring the role of neuroendocrine cells in obesity (Supplementary Fig. 6c).

To elucidate systemic obesity-associated alterations, we performed comprehensive analysis of DEGs (Supplementary Table 7). KEGG pathway analysis demonstrated that DEGs in glandular and neural tissues were enriched for hormone signaling and synaptic function (Fig. 6c). Metabolic profiling of obese mice revealed profound alterations in glucose and lipid metabolism, characterized by upregulated gluconeogenesis and fatty acid synthesis. Notably, amino acid metabolism was also substantially remodelled, with elevated branched-chain amino acids (BCAAs) metabolism^34^ but decreased arginine and aspartate metabolism involved in the urea cycle^35^, potentially contributing to hepatic toxin accumulation and oxidative stress. These transcriptional changes were strongly corroborated by our metabolomics data (Supplementary Fig. 7). AMPK signaling, a key metabolic regulatory pathway, along with pathways governing cell growth and differentiation, were broadly downregulated in obese mice.

TFs regulatory network revealed tissue-specific metabolic dysregulation (Fig. 6d). Notably, Tcf7l2^36^ showed divergent expression – upregulated in kidney but downregulated in liver and pituitary, suggesting distinct tissue adaptations that affect glucose metabolism and hormonal regulation^37^. The metabolic regulators FOXO1/3^38^ were downregulated in liver and colon, while widespread THRB downregulation indicated impaired thyroid hormone signaling. In summary, we present the first comprehensive multi-tissue single-nucleus transcriptomic atlas of obesity, revealing both conserved and tissue-specific transcriptional changes across multiple systems.

## Discussion

In this study, we present UURNA-seq, an ultra-throughput and ultra-sensitivity platform for single-nucleus total RNA sequencing. Four levels of indexing significantly enhance the throughput of single-cell RNA assays. In contrast to conventional split-pool protocols^7^, UURNA-seq employs direct double-strand extension after hybridization and therefore requires no ligation. Direct PCR from the cell pellet maximizes cDNA release, making UURNA-seq the most sensitive method among high-throughput strategies. Random priming further improves the sensitivity of nucleus-based analysis and reduces the requirements on sample quality. Single-nucleus sequencing is compatible with a broader range of samples, particularly for limited clinical specimens-a significant advantage that addresses a major limitation in the wide adoption of current full-length sequencing technologies^13,15,23^. By capturing total RNA in nuclei, UURNA-seq gives access to non-coding RNAs, particularly lncRNAs, and sncRNAs.

Using UURNA-seq, we constructed a comprehensive mouse brain atlas that successfully identified distinct neuronal populations across brain regions without precise anatomical dissection, demonstrating the high specificity of the methodology. Through comparison with 10× Genomics datasets, we elucidated the unique advantages of UURNA-seq, particularly its superior capture efficiency for non-coding RNAs and transcription factors.

Despite these advantages in throughput and sensitivity, UURNA-seq is currently limited by sequencing cost and computational demand. Advances in next-generation sequencing technologies and computational algorithms, such as more efficient data compression and cloud-based pipelines, are anticipated to address the challenges posed by the growing scale of single-cell genomic data. We used fixed nuclei for random primer-mediated reverse transcription and found that the optimal fixation protocols may vary across different cell types. Efforts to develop improved fixation reagents for in-cell biochemical reactions, which are currently lacking in the field, are crucial. Additionally, the efficiency of single-nucleus isolation varies among different tissue types, and while our comparison of detergents provides a starting point for tissue-specific optimization, a more systematic screening across diverse tissue types will be necessary. Furthermore, while profiling total RNA in single nuclei provides a good representation of transcriptional activity, our measurements may differ from single-cell profiles at the mRNA and non-coding RNA level. Nevertheless, these limitations do not diminish the broad applicability of UURNA-seq; rather, they mark clear directions for future refinement as the platform is applied to increasingly diverse tissue types and clinical contexts.

## Materials and methods

### Oligonucleotides

The sequences and modifications for all oligonucleotides are listed in Supplementary Table 1.

### Cell culture

HEK 293T and NIH/3T3 cells were passaged every 48 hours and maintained in a humidified incubator at 37°C with 5% CO□. Cells were cultured in Dulbecco’s Modified Eagle Medium (DMEM, Gibco) supplemented with 10% fetal bovine serum (FBS, Gibco) and 1% penicillin–streptomycin (Gibco). For passaging, cells were rinsed twice with 2 mL of 1× DPBS followed by dissociation with 1 mL of 0.5% Trypsin for 3-5 minutes at 37°C. Trypsinization was terminated by adding 2 mL of DMEM containing 10% FBS. Cells were subsequently centrifuged at 300g for 5 minutes. After aspirating and discarding the supernatant, cell pellets were washed twice with 1× DPBS before nucleus preparation.

### Mice

Wild-type C57BL/6J mice were housed in groups of three to five per cage in a Specific Pathogen Free (SPF) facility with ad libitum access to food and water. The housing environment was maintained at a controlled temperature (20-22°C), humidity (30-70%), and under a standardized 12-hour light-dark cycle. Adult female mice (8-13 weeks of age) and prepubertal mice (3 weeks of age) were used for the experiments. All animal procedures were performed under protocols approved by the Animal Ethics Committee and complied with all relevant regulatory standards.

### Nucleus isolation for cell lines and primary mouse tissues

#### Cell lines

After trypsin digestion, cells were washed twice with 1× DPBS. Cell pellets were resuspended in ice-cold nucleus isolation buffer (NIB: 10 mM Tris-HCl pH 7.5, 10 mM NaCl, 3 mM MgCl□, 0.1% IGEPAL CA-630, 0.1% Tween-20, 1 mM dithiothreitol (DTT), and 0.4 U/µL murine RNase inhibitor) and incubated on ice for 5 minutes. The suspension was subsequently diluted with nucleus wash buffer (NWB: 10 mM Tris-HCl pH 7.5, 10 mM NaCl, 3 mM MgCl□, and 0.1% Tween-20) and centrifuged at 500g for 5 minutes at 4°C. For species mixing experiments, nuclei were combined in equal proportions at this stage based on appropriate concentration calculations. Following two additional washes, the nuclei pellet was resuspended in 100 μL of 1× DPBS containing 0.4 U/μL murine RNase inhibitor. Nuclear fixation was performed by adding 5 mL of pre-chilled RNase-free 4% PFA fixation buffer, followed by incubation on ice for 1 hour with occasional gentle mixing. The fixation reaction was quenched by adding 750 μL of 2.5 M glycine and incubating on ice for 15 minutes. Fixed nuclei were then washed twice with NWB supplemented with 0.4 U/μL murine RNase inhibitor and filtered through a 40 μm cell strainer.

#### Primary tissues

Mice were anesthetized with 1% sodium pentobarbital, and the intestine was removed prior to perfusion with 1× DPBS. Other target tissues were excised and rinsed with DMEM. After blotting tissues dry with absorbent paper, they were pulverized in liquid nitrogen using a stainless-steel blender. The resulting tissue powder was immediately homogenized in ice-cold NIB buffer. After a 3-minute incubation, the homogenate was transferred to 5 mL of NWB and filtered through a 40 μm cell strainer. The nuclei suspension was then centrifuged at 500g for 5 minutes at 4°C.

While this extraction protocol was suitable for most tissues, a gentler approach was implemented for more fragile tissues to obtain high-quality nuclei. Briefly, these tissues were homogenized in a 3 mL glass Dounce homogenizer with 5-10 strokes of a loose pestle (A) followed by 5-10 strokes of a tight pestle (B) in homogenization buffer^39^ (250 mM sucrose, 25 mM KCl, 5 mM MgCl_2_, 10 mM Tris-HCl, 1 mM DTT, 0.4 U/μL Murine RNase Inhibitor and 0.1% Triton X-100 in nuclease-free water). The homogenate was filtered through a 40 μm cell strainer and centrifuged at 500g for 5 minutes at 4°C. After removing the supernatant, the pellet was resuspended in homogenization buffer without Triton X-100. A tissue-specific nucleus isolation protocol is detailed in Supplementary Table 2.

After washing, nuclei from primary tissues were fixed using the same procedure as for cell lines, but with a reduced fixation time of 20 minutes. For tissues yielding few nuclei, fixation was performed in 1.5 mL centrifuge tubes to minimize loss due to cell adhesion. After fixation, nuclei were washed with the corresponding buffer and filtered again using either a 40 μm or 20 μm strainer. The final nuclei pellet was resuspended in 1× PBS containing 0.4 U/μL murine RNase inhibitor and adjusted to the desired concentration.

### UURNA-seq procedure

#### In situ reverse transcription

The first round of barcoding was performed by in situ reverse transcription (RT). Prior to processing the nuclei suspension, the RT reaction mix was prepared. RT primers (2 µL) were aliquoted into a 96-well plate, incubated at 95°C for 3 minutes in a thermal cycler, then immediately transferred to ice. The RT mix for each well comprised: 2 µL 5× RT buffer (Thermo Scientific), 2 µL 40% PEG-8000, 0.5 µL 5% Triton X-100, 0.5 µL 10 mM dNTP mix (Thermo Scientific), 0.5 µL Maxima H Minus (Thermo Scientific), and 0.5 µL RNase inhibitor (Vazyme Biotech). This RT mix was dispensed into the primer-containing wells and mixed by centrifugation at 3,000g for 5 minutes. Subsequently, 2 µL of nuclei suspension (∼50,000 nuclei) was added to each well. Ten cycles of multiple annealing were performed according to the following temperature profile: 8°C for 12 seconds, 15°C for 30 seconds, 20°C for 45 seconds, 25°C for 1 minute, 30°C for 1 minute, and 42°C for 2 minutes. The plate was then incubated at 42°C for 30 minutes with rotation to facilitate an efficient RT reaction. The reaction was terminated by adding 10 µL of 40 mM EDTA to each well and incubating at 37°C.

#### Exonuclease □ treatment and poly(A) tailing

After reaction termination, the nuclei were pooled and washed three times with PBST (0.05% Tween-20 in 1× DPBS). Residual primers were removed by Exonuclease I treatment. Nuclei were resuspended in 50 µL of Exonuclease I mix per million cells, consisting of 1× Exonuclease I buffer and 50 U Exonuclease I (NEB). The mixture was incubated at 37°C for 30 minutes with continuous mixing on a rotary mixer. The reaction was terminated by adding an equal volume of 40 mM EDTA. The nuclei were then washed three times prior to the next reaction. For poly(A) tailing, nuclei were resuspended in 50 µL of Terminal transferase mix per million cells, comprising 1× terminal transferase buffer (Roche), 5 mM CoCl□, 100 µM dATP, and 200 U terminal transferase (Roche). The reaction was performed at 37°C for 15 minutes with rotation and quenched by adding an equal volume of 40 mM EDTA.

#### Hybridization and block

After three washes with PBST, nuclei were resuspended in hybridization mix (per well: 0.5 µL 10× hybridization buffer [prepared in ddH_2_O containing 100 mM Tris-HCl (pH 8.0), 500 mM NaCl, and 10 mM DTT], 0.05 µL 10% Triton X-100, 0.5 µL 50% PEG-8000, and 1.95 µL ddH□O). The suspension was distributed across 96-well plates, and 2 µL of 10 µM beads linker 1.2a primer was added to each well for second-round barcoding. Hybridization proceeded with 10-minute rotation at 37°C, followed by 20 minutes at room temperature. To sequester excess hybridization primers, 2 µL of blocking primer (20 mM) was added, and plates were rotated at room temperature for 15 minutes. After blocking, nuclei were pooled in wash buffer, centrifuged at 500g for 5 minutes at 4°C, and washed twice with PBST.

#### Extension

Nuclei were resuspended in extension buffer (per well: 0.25 µL 10× Blue buffer [prepared in ddH□O containing 100 mM Tris-HCl (pH 8.0), 500 mM NaCl, 100 mM MgCl□, and 10 mM DTT], 0.5 µL 10 mM dNTP mix, and 1.75 µL ddH□O) and diluted according to the number of barcode combinations in the first two rounds. After distribution across 96-well plates, samples underwent two freeze-thaw cycles at –80°C. An enzyme mix (2.5 µL containing 0.25 µL 10× Blue buffer, 0.125 µL Klenow, 0.125 µL exonuclease I, 0.025 µL RNase H, and 1.975 µL ddH□O) was added to each well. The extension reaction was performed at room temperature for 30 minutes, followed by a thermal cycling program: 37°C for 10 minutes, 40°C for 5 minutes, 45°C for 5 minutes, 50°C for 5 minutes, then held at 4°C.

#### Lysis and PCR preamplification

A lysis buffer (1 µL; prepared as 12 µL 10% SDS, 0.5 µL Proteinase K (Sangon Biotech, 20 mg/mL), and 87.5 µL ddH□O per 100 µL) was added to each well and incubated at 55°C for 15 minutes. SDS was neutralized by adding 6 µL of 10% Tween-20. PCR mix (15 µL containing 14 µL HiFi HotStart ReadyMix (Kapa Biosystems) and 1 µL of 2.5 µM ATAC R1.1 primer) was added, followed by 1 µL of 2.5 µM beads linker 1.2b primer containing the third barcode. The PCR thermal cycling program consisted of: 72°C for 5 minutes, 95°C for 3 minutes; two cycles of 98°C for 20 seconds, 65°C for 1 minute, and 72°C for 1 minute; four cycles of 98°C for 20 seconds, 65°C for 30 seconds, and 72°C for 1 minute; 72°C for 5 minutes; and a final hold at 4°C.

#### Purification and library preparation

Eight-well pooled PCR products were purified using 1.2× VAHTS DNA Clean Beads (Vazyme Biotech). The purified product (23 µL in DEPC-treated water) was added to a PCR mix consisting of 25 µL HiFi HotStart ReadyMix, 1 µL 10 µM P5-ATAC primer, and 1 µL 10 µM i7-ATAC primer. The PCR program comprised: 95°C for 3 minutes; two cycles of 98°C for 20 seconds, 65°C for 1 minute, and 72°C for 1 minute; seven to ten cycles of 98°C for 20 seconds, 65°C for 30 seconds, and 72°C for 1 minute; 72°C for 5 minutes; followed by a hold at 4°C. Libraries were size-selected (250-700 bp) using two rounds of VAHTS DNA Clean Beads purification and quantified using a Qubit fluorometer (Invitrogen).

### Sequencing of UURNA-seq library

The purified double-stranded DNA library was denatured at 98°C and subsequently circularized into a single-stranded DNA (ssDNA) library using a VAHTS® Circularization Kit for MGI (Vazyme Biotech). The ssDNA library was then amplified using a DNBSEQ DNB preparation kit (MGI). The resulting amplified DNA nanoballs (DNBs) were sequenced with TrueSeq sequencing primers on an MGI DNBSEQ-T7 platform.

### Preprocessing of UURNA-seq data

The raw FASTQ-format sequencing data from a DNBSEQ-T7 were first split into i7-indexed sub-libraries using splitBarcode (v.0.1.6) [https://github.com/MGl-tech-bioinformatics/splitBarcode]. We preprocessed the sub-library data with the Drop-seq^28^ core computational tool. Using the “TagBamWithReadSequenceExtended” function, we extracted the 20-nt barcode combinations from hybridization and the first PCR in Read 1, together with the 10-nt RT barcodes and 7-nt UMI in Read 2. We then corrected the cell barcodes for each of the three rounds of split-pooling with a custom Python script, allowing a Hamming distance of ≤1 in each round. The potential poly(A) tail on Read 2 was trimmed with cutadapt^40^ (v.3.5), applying the read-length filter: –-minimum-length=20. Next, we mapped reads using STAR^41^ (v.2.5.1) with default parameters. We aligned reads from 3T3 cells and 293T cells to a merged hg19-mm10 genome reference (provided by Drop-seq group; GSE63269), and reads from WT mice to the Mus_musculus.GRCm38.88 genome. For quality control, we filtered out cells with fewer than 500 detected transcripts, along with cells in which mitochondria-encoded genes contributed a high proportion of transcript counts (>∼5–20%, depending on the dataset). Finally, we used scDblFinder (v.1.16.0)^42^ to detect potential doublets in two rounds. Setting the samples variable to “index” for the first round and “sample” for the second, we run the “scDblFinder” function to detect and remove potential doublets within each index or each mouse.

### Dimensionality reduction and clustering

We employed the Seurat package (v.4.4.0) for dimensionality reduction and cell clustering for both the tissue and integrated datasets. The data were log2(TP10K+1)-transformed, with the number of UMIs and the percentage of mitochondrial gene content regressed out. We selected 2,000 genes with the “FindVariableFeatures” function (“vst” method) as inputs for initial principal component analysis (PCA). The number of principal components (PCs) used for nonlinear dimensionality reduction (UMAP)^43^ ranged from 15 to 40. For clustering, we varied the resolution parameters from 0.5 to 2 in the “FindClusters” function. These parameters, including the number of PCs and the resolution, were tailored to each tissue sample.

### Benchmarking current sc/snRNA methods

#### Comparison with cell line data

We benchmarked against Smart-seq3 (<u>E MTAB-8735</u>), FLASH-seq (Sequence Read Archive, <u>PRJNA816486</u>), sci-RNA-seq (<u>GSM2599699</u>), SPLiT-seq (<u>GSM3017263</u>) and 10× Genomics (https://www.10xgenomics.com/datasets/10-k-1-1-mixture-of-human-hek-293-t-and-mouse-nih-3-t-3-cells-3-v-3-1-chromium-controller-3-1-standard-6-1-0). We trimmed homopolymers, extracted cell barcodes and UMIs, then mapped all raw cell line reads to the combined hg38-mm10 genome with STAR (v2.5.2) using default parameters. For gene detection, only cells with the highest read counts were retained (30,000 reads for the saturation curve). We down-sampled the aligned BAM files and analyzed only reads mapping to coding and intronic regions. Average gene body coverage was calculated with QoRTs^44^ (v.1.3.6). To classify transcript biotypes, we annotated aligned reads with TagReadWithGeneFunction from Drop-seq tools against the gene features defined in the GTF annotation. Reads mapping to more than one gene locus were defined as ‘Multi-mapped’.

### Comparison between 10× Chromium and UURNA-seq brain data

For scRNA-seq data, we benchmarked UURNA-seq against 10× Chromium 3′ v1/2 platform using the mouse nervous system data. We selected the same regions profiled by UURNA-seq and downloaded them from https://www.ncbi.nlm.nih.gov/sra/SRP135960. We utilized MetaNeighbor (v.1.22.0)^45^ to retain cells with a correlation > 0.95 for further analysis. Differentially expressed genes (DEGs) within the filtered cell types of the two methods were identified using the “FindMarkers” function in Seurat (|log2FC| > 1; adjusted *P* < 0.01). GO enrichment analysis of the DEGs was performed by clusterProfiler (v.4.10.0)^46^ R package. To compare tissue dissociation-included stress, we applied a published dissociation signature^47^ to the snRNA-seq and scRNA-seq profiles using the “AddModuleScore” function in Seurat. The score was computed based on normalized log-transformed gene expression values (log2(TP10K+1)) for each cell type shared by the two datasets.

We correlated pseudobulk expression profiles between cells and nuclei using Spearman correlation of log2-transformed values for protein-coding genes. Residuals were computed by fitting a linear model (“resid (lm (cells-nuclei ∼ 0))”) to the scRNA-seq and snRNA-seq expression profiles. Genes with residuals above the 97.5th or below the 2.5th percentile were classified as divergent. Poly-A content was quantified by identifying stretches of at least 20 consecutive adenines using the BSgenome.Mmusculus.UCSC.mm10 package; total poly-A length per gene was calculated as the sum of these adenine units, and gene length was taken from the same package.

To compare snRNA-seq data between 10× Genomics and UURNA-seq, we obtained mouse cortex (<u>GSE140511</u>) and cerebellum (<u>GSE165371</u>) datasets and selected nuclei from the same regions profiled by UURNA-seq. Subsequently, only nuclei with correlation > 0.995 in the cerebellum and > 0.85 in the cortex were retained for further analysis. For down-sampling and comparison of gene numbers, cerebellar nuclei with at least 1,000 sequenced reads in both coding and intronic regions were selected. Gene biotypes were annotated from the Ensembl GTF (version 88) for the mm10 genome.

### Analysis of lncRNAs

Unsupervised clustering of lncRNAs followed the same pipeline as for total RNA, with the exception that doublet identification and removal were omitted because only nuclei passing the ‘singlet’ filter were retained. We used the method of Upsilon^48^ to measure the cell type specificity scores for protein-coding genes and lncRNAs.

### Construction of single-cell gene co-expression networks

Following the methods of Morabito et al., we constructed co-expression networks from snRNA-seq data of the cerebellum using hdWGCNA (v.0.4.4)^49^. To ensure analytical quality, we retained only genes expressed in at least 5% of cells. Metacell transcriptional profiles were built for each cell type using the MetacellsByGroups function by aggregating 25 cells per metacell. Based on parameter scanning results from the TestSoftPowers function, we determined a soft-power threshold of β = 5. Subsequently, we constructed the co-expression network using the ConstructNetwork function with the following parameters: networkType = “signed”, TOMType = “signed”, soft_power = 5, deepSplit = 4, detectCutHeight = 0.995, minModuleSize = 50 and mergeCutHeight = 0.2. Module eigengenes (MEs) were assessed using the ModuleEigengenes function, and the connectivity of each gene to the module eigengenes was computed using the ModuleConnectivity function. The RunModuleUMAP function embedded the network structure in two dimensions for visualization. Finally, functional enrichment analysis of genes within each module was performed using the enrichR^50^ (v.3.4) tool, querying GO_Biological_Process_2023, GO_Cellular_Component_2023, and GO_Molecular_Function_2023. For co-expression analysis of the stromal lineage datasets, we employed the same approach, except that each metacell aggregated 30 cells and the soft-power threshold was adjusted to 9. Distributions of MEs were compared in WT mice for each cell type using a two-sided Wilcoxon rank-sum test (R function wilcox.test).

## Data availability

Raw data generated in this study is available in Gene Expression Omnibus under accession number (<u>GSE287434</u>). Previously published datasets used in this study are available under the following accession codes: Smart-seq3 (<u>E MTAB-8735</u>), FLASH-seq (Sequence Read Archive, <u>PRJNA816486</u>), sci-RNA-seq <u>(GSM2599699</u>), SPLiT-seq (<u>GSM3017263</u>), 10× Genomics <u>(10× Genomics official website).</u> 10× Genomics scRNA-seq data of mouse whole brain (<u>GSM6617915</u>), 10× Genomics snRNA-seq of mouse cortex (<u>GSE140511</u>), 10× Genomics snRNA-seq of mouse cerebellum (<u>GSE165371</u>). GRO-seq of cortical neurons (<u>GSM6626513</u>).

## Code availability

Code and analysis scripts related to the methods are available in the GitHub repository (https://github.com/LifengMa/UUseq)

## Figure

**Figure S1.**
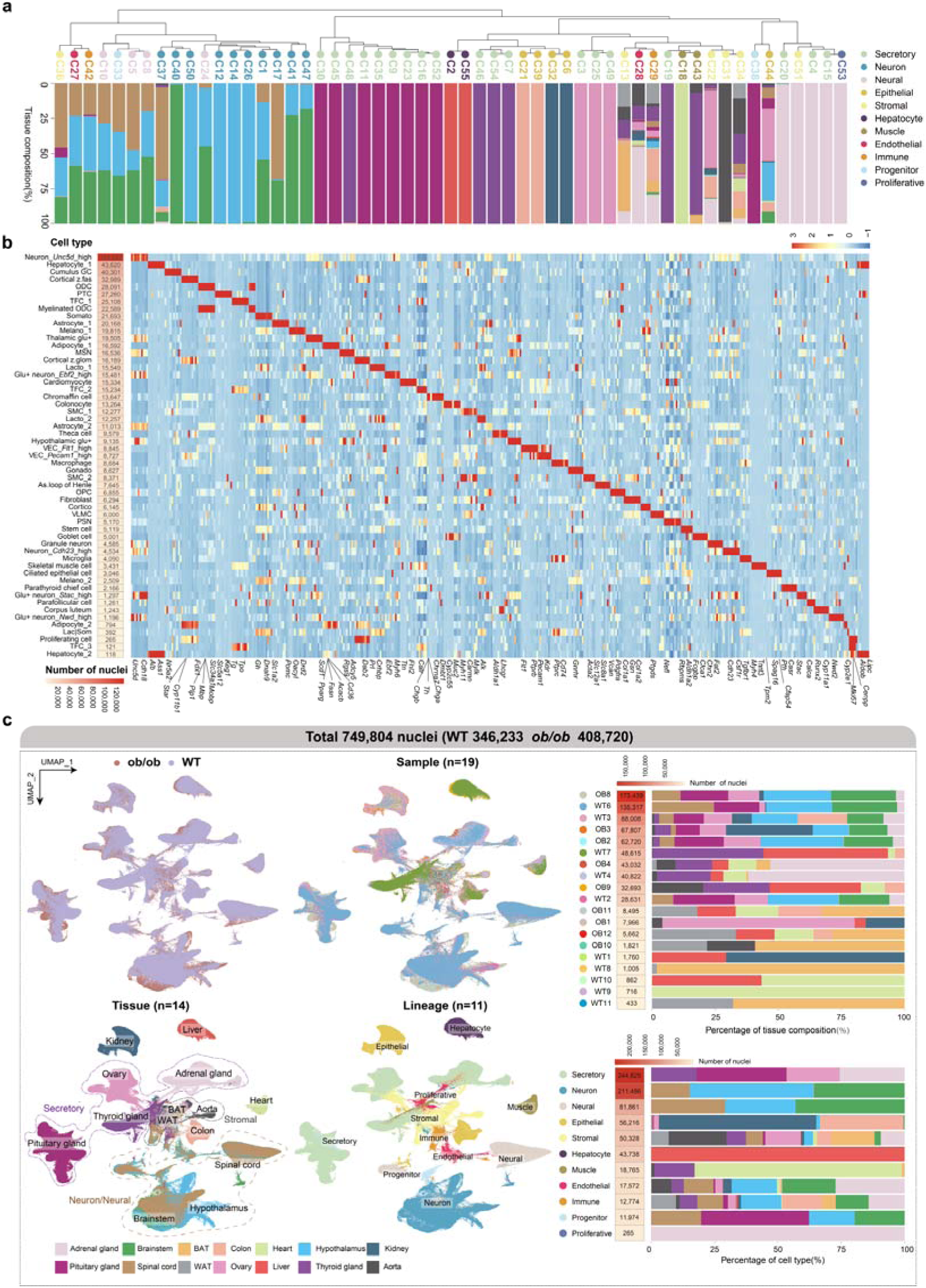
Characterization of the cross-tissue snRNA-seq atlas. **a**, Hierarchical clustering of tissue composition across various cell clusters. The dendrogram at the top shows the clustering of the cell types by expression profile. Each column corresponds to a cell type and indicates the proportion of that cell type within each tissue. **b,** Heatmap of differentially expressed genes across annotated cell types. Nuclei counts for each cell type are shown on the left. Significantly differentially expressed genes are highlighted below. **c,** UMAP visualization of 749,804 nuclei (WT 346,233, *ob/ob* 408,720), color-coded by genotype, sample (n = 19), tissue (n=14) and lineage (n=11). Bar charts of the nuclei number and the percentage of tissue composition for each sample (top) and lineage (bottom).

**Figure S2.**
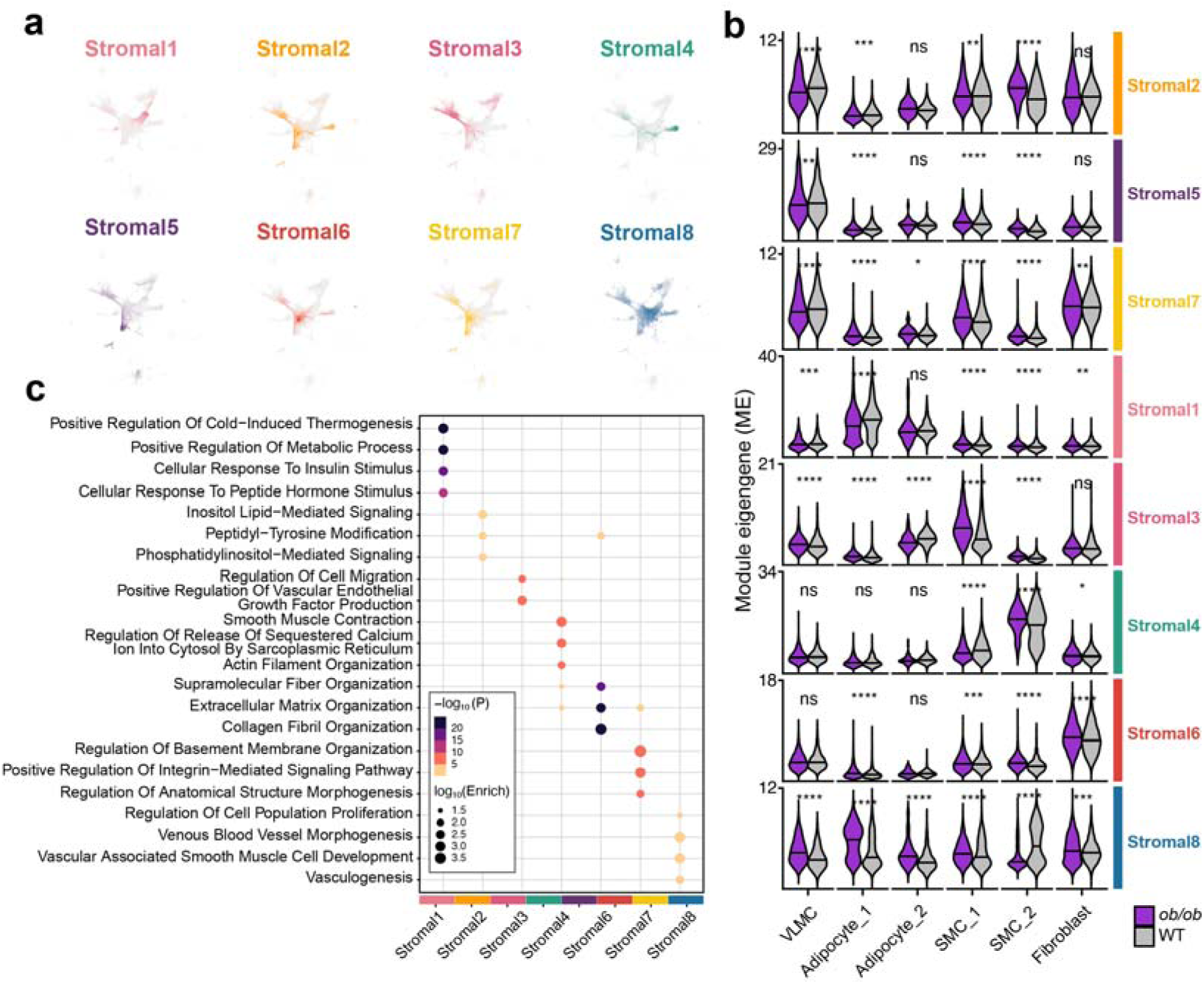
Co-expression network analysis of stromal lineage. **a**, snRNA-seq UMAP plots as in stromal lineage, colored by MEs for co-expression modules. **b,** Violin plots showing MEs in each stromal cluster. Two-sided Wilcoxon test was used to compare *ob/ob* versus WT samples. ns, not significant, *P* > 0.05; *, *P* < 0:05; **, *P* < 0:01; ***, *P* < 0:001; ****, *P* < 0:0001. **c,** Selected GO enrichment results for each co-expression module.

**Figure S3.**
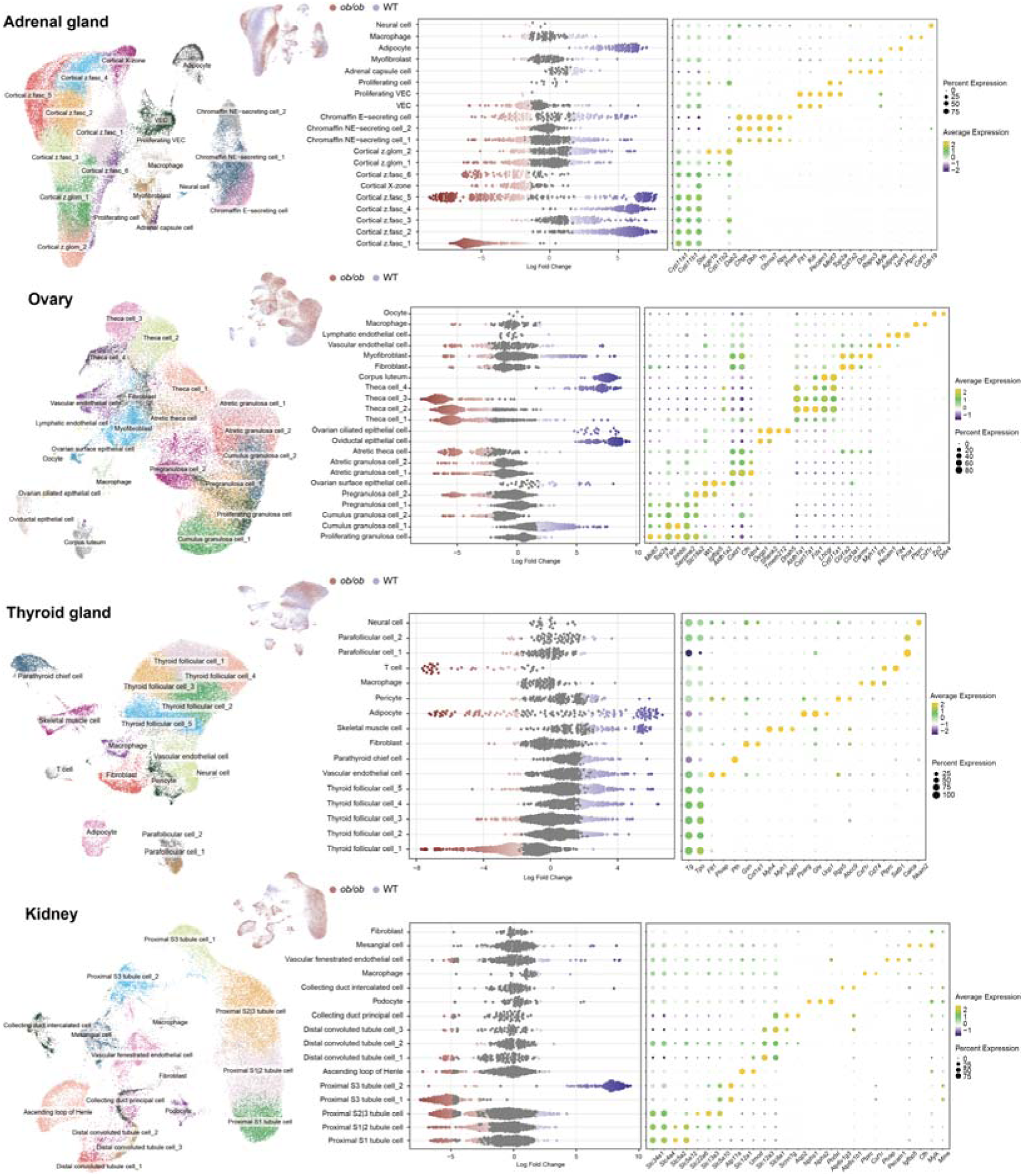
Representative tissue-level UMAP visualization of UURNA-seq datasets. UMAP visualization of single-nucleus data from adrenal gland, ovary, thyroid gland and kidney, color-coded by cell-type cluster (left). Beeswarm plot of the distribution of log fold change between lean and obese mice across neighborhoods containing cells from different cell type clusters (middle). Differentially abundant neighborhoods at 10% FDR are colored. Dot plots of cell-type-specific marker gene expression (right). Dot size corresponds to the fraction of cells expressing the gene and the color gradient represents the average expression level within each cell type.

**Figure S4.**
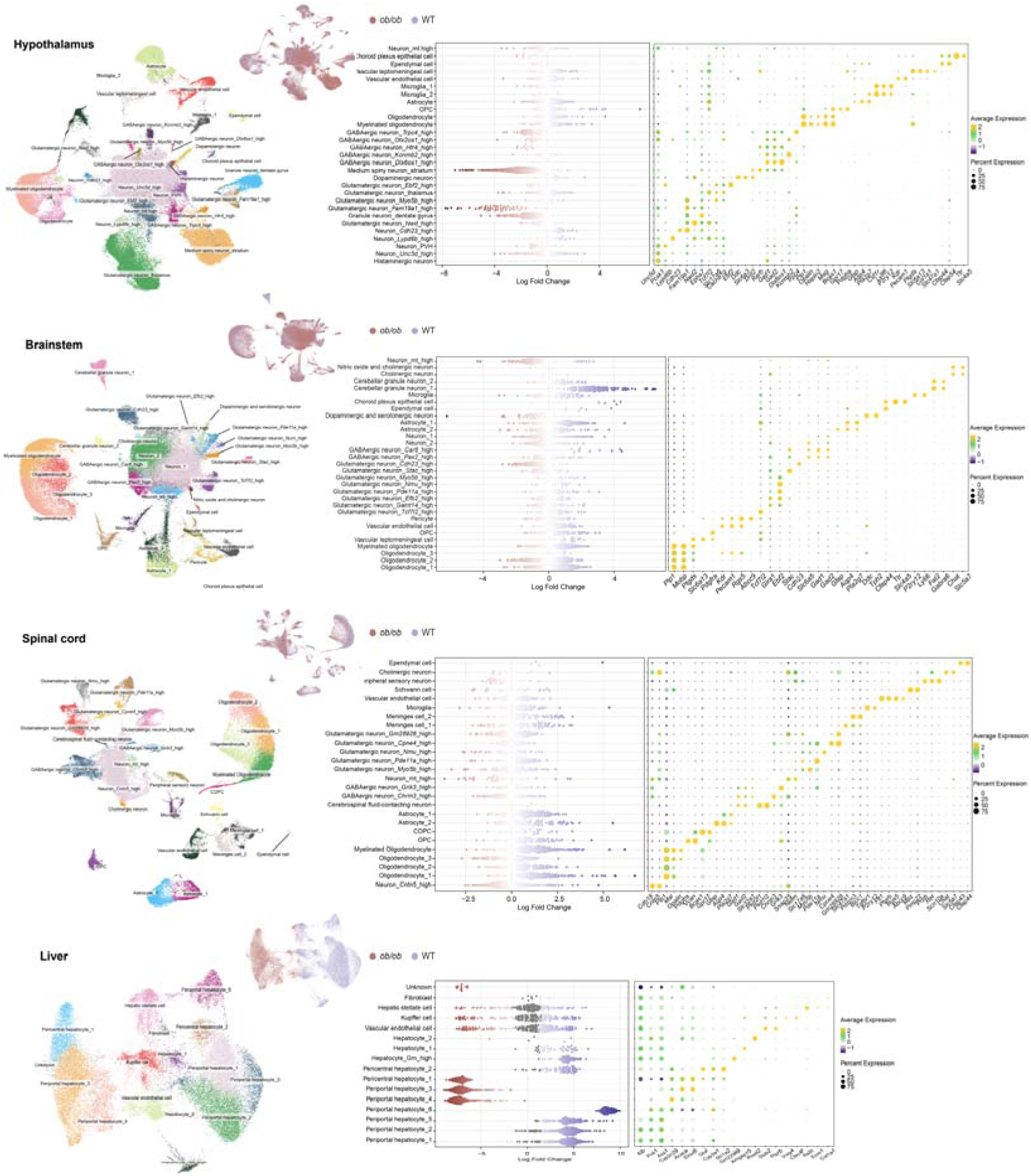
Representative tissue-level UMAP visualization of UURNA-seq datasets. UMAP visualization of single-nucleus data from hypothalamus, brainstem, spinal cord and liver, color-coded by cell-type cluster (left). Beeswarm plot of the distribution of log fold change between lean and obese mice across neighborhoods containing cells from different cell type clusters (middle). Differentially abundant neighborhoods at 10% FDR are colored. Dot plots of cell-type-specific marker gene expression (right). Dot size corresponds to the fraction of cells expressing the gene and the color gradient represents the average expression level within each cell type.

**Figure S5.**
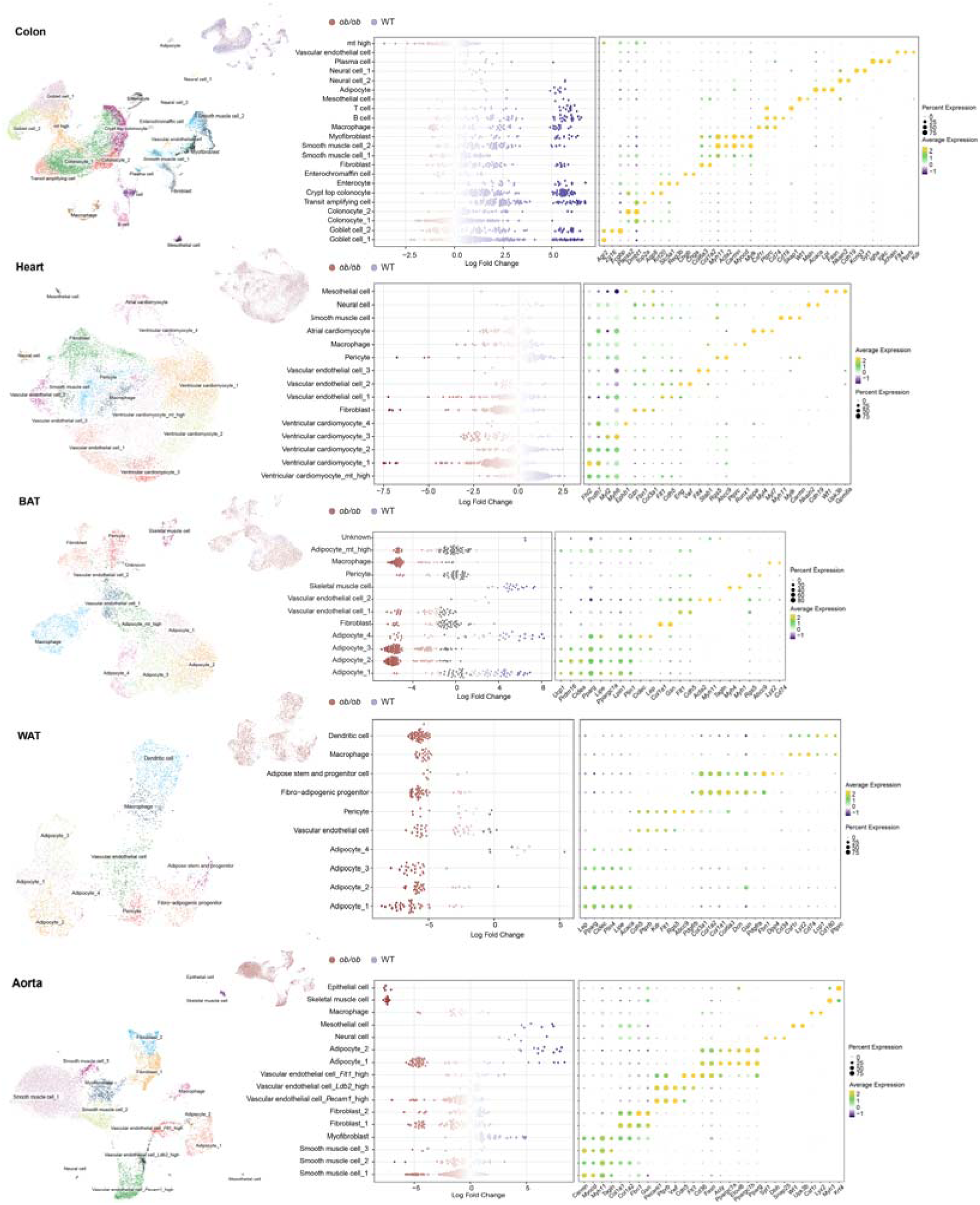
Representative tissue-level UMAP visualization of UURNA-seq datasets. UMAP visualization of single-nucleus data from colon, heart, brown adipose tissue (BAT), white adipose tissue (WAT) and aorta, color-coded by cell-type cluster (left). Beeswarm plot of the distribution of log fold change between lean and obese mice across neighborhoods containing cells from different cell type clusters (middle). Differentially abundant neighborhoods at 10% FDR are colored. Dot plots of cell-type-specific marker gene expression (right). Dot size corresponds to the fraction of cells expressing the gene and the color gradient represents the average expression level within each cell type.

**Figure S6.**
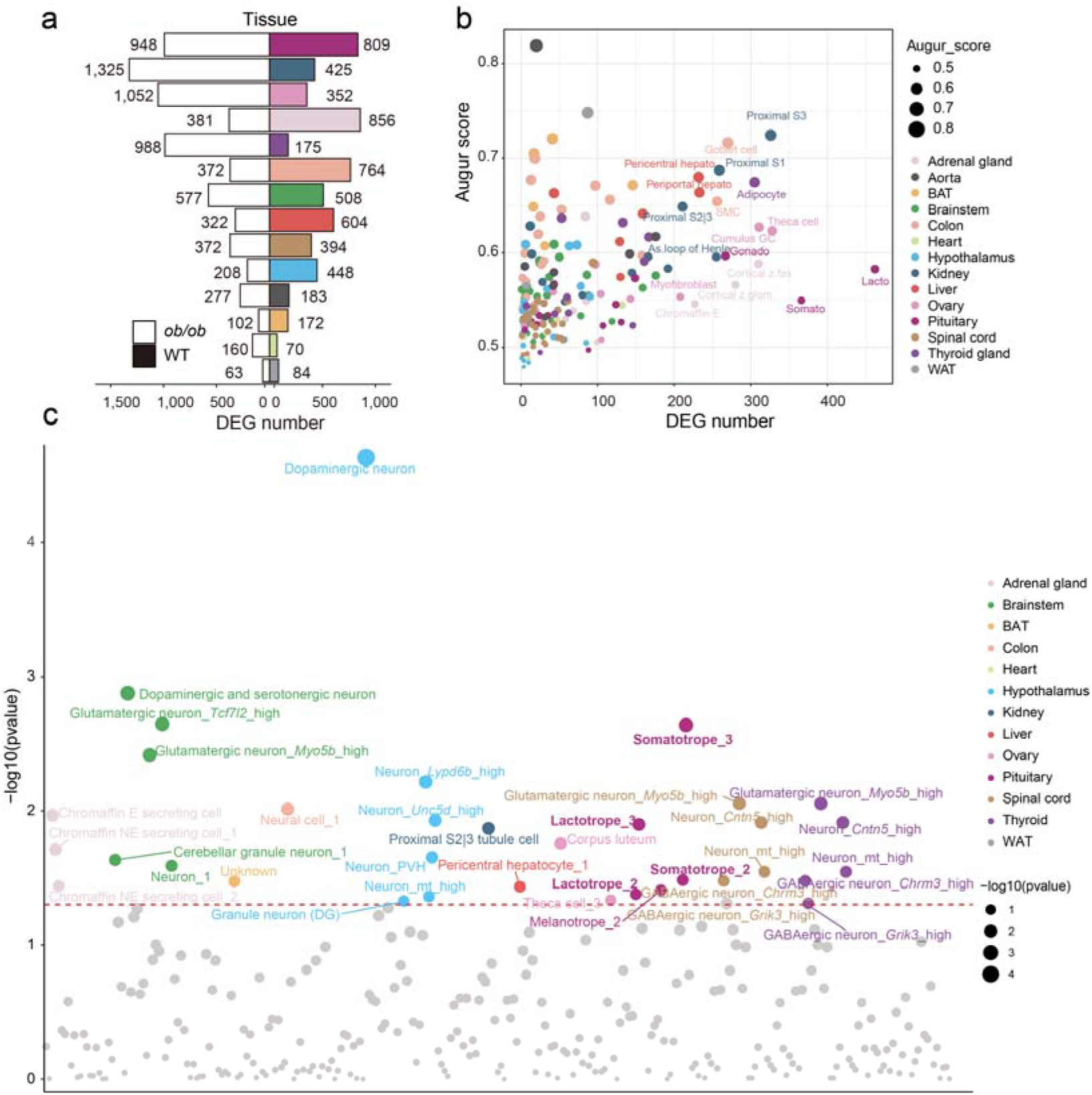
Overview of DEGs and their association with cell types and GWAS data. **a**, Bar plots of DEG numbers in *ob/ob* (left) and WT (right). **b,** Bubble plot of DEG number and Augur score per cell type. Dot size indicates the Augur score and dot color indicates the tissue. **c,** CELLLECT *P* values for the association between cell types of tissues in the UU-RNA-seq dataset and the GWAS BMI trait. The red line indicates the significance threshold of *P* = 0.05.

**Figure S7.**
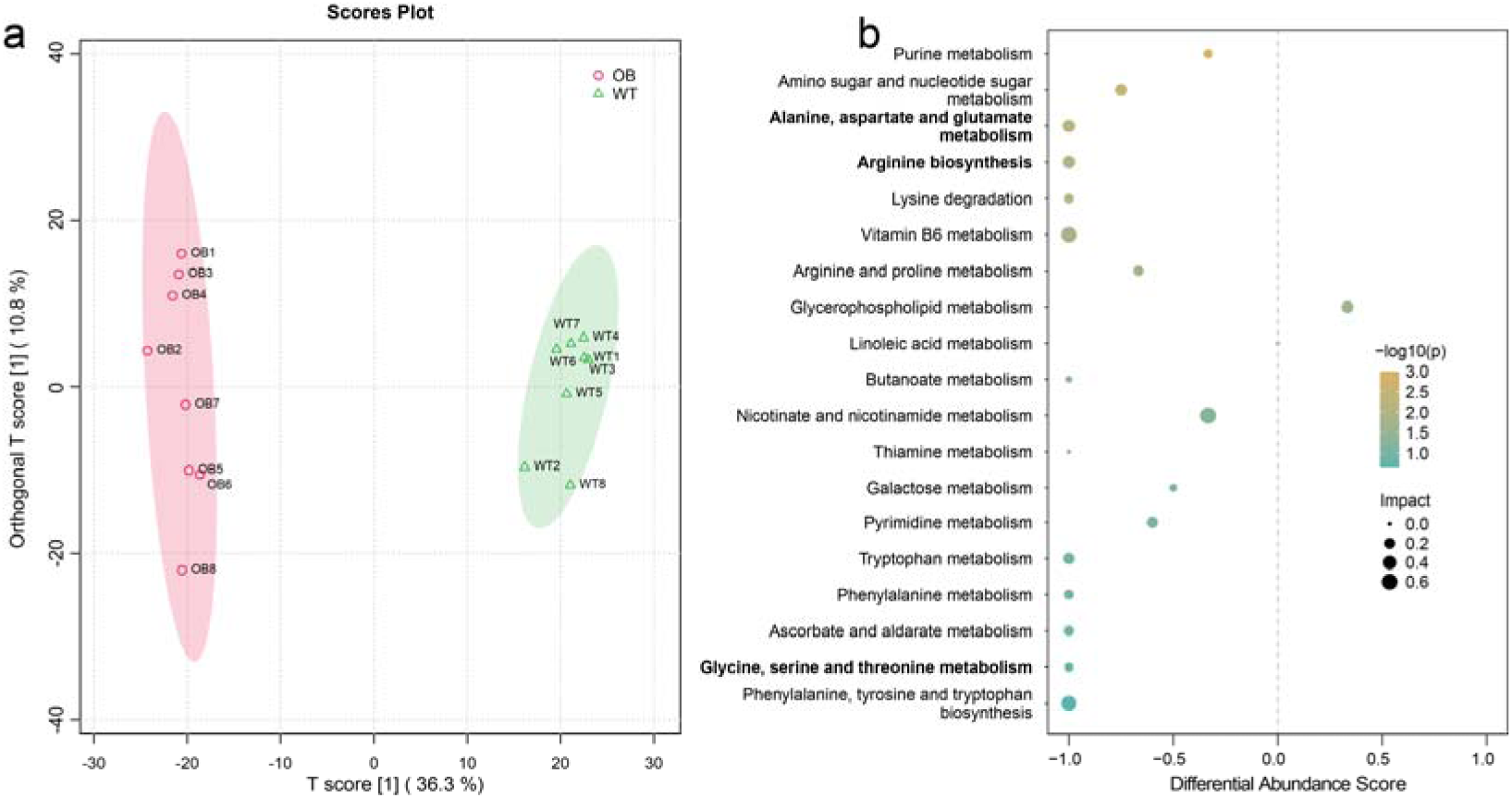
Alterations in the hepatic metabolome. **a**, Scores plot from orthogonal partial least squares discriminant analysis (OPLS-DA) showing distinct metabolic profiles for *ob/ob* (circles) and WT (triangles) mice. The T score represents the prediction strength and the orthogonal T score indicates the orthogonal distance. Ellipses represent the 95% confidence intervals for the two groups. **b,** Bubble plot of the differential abundance of metabolic pathways between the two groups. The x axis represents the differential abundance score. Pathways with FDR ≤ 0.05 are highlighted in color, with the color intensity representing the statistical significance (-log10(*P*)) and the bubble size corresponding to the pathway impact.

## Notes

https://www.ncbi.nlm.nih.gov/geo/query/acc.cgi?acc=GSE287434

### Competing Interest Statement

The authors have declared no competing interest.

### Funding

- National Natural Science Foundation of China: 32570678, 32271491, 32330061
- Zhejiang Province Vanguard Goose-Leading Initiative: 2025C01114
- Funds for Creative Research Groups of China: T2121004

